# Screening of associated bacteria of *Alexandrium catenella* on affecting PSTs production and influence mechanism

**DOI:** 10.1101/2024.01.31.578139

**Authors:** Shanmei Zou, Xinke Yu, Tiantian Sun, Lina Wei, Xuemin Wu

## Abstract

*Alexandrium* causes serious food safety and human death due to paralytic shellfish toxins (PSTs) production. The associated bacteria can affect PSTs production of *Alexandrium*. However, the influencing mechanism is still unclear. Here we firstly screened functional associated bacteria for affecting PSTs production of *Alexandrium catenella* in Yangtze Estuary and further studied their influence on physiological process and molecular regulation of *A. catenella*. Thirteen bacteria strains for affecting PSTs production of *A. catenella* were selected. The *A. catenella* strains co-cultured with different functional associated bacteria all produced more PSTs than axenic strain with antibiotic treatment. Compared with axenic *A. catenella*, the non-axenic *A. catenella* produced more algal cells, soluble sugar, soluble protein and neutral lipid. By RNA-seq, it was found that non-axenic *A. catenella* produced more upregulated functional genes than axenic *A. catenella*. The biosynthesis of cofactors and spliceosome were the dominant different pathways between axenic and non-axenic *A. catenella* strains. The sxtA expression was closely related with Arginine and proline metabolism, Arginine biosynthesis, Fatty acid biosynthesis, TCA cycle and Glutathione metabolism, which were all downregulated in axenic *A. catenella*. Meantime, the non-axenic *A. catenella* under nitrogen deprivation produced less PSTs and functional genes than non-axenic strain under common culture condition, indicating the nitrogen significance for PSTs production. The detailed signal molecular released by associated bacteria for regulating PSTs of *A. catenella* needs to be further studied.

## 1. Introduction

Harmful algal blooms (HABs) occur globally and have produced severe effects on fisheries, aquaculture, tourism and recreation[1, 2]. Dinoflagellates, a large group of phytoplanktonic organisms during marine HABs, play a crucial role in food webs and in balancing ecosystem energy fluxes [3–8]. Many marine dinoflagellates can cause food safety and human fatalities through their production of paralytic shellfish toxins (PSTs) which accumulate in shellfish through the food chain when contaminated seafood is consumed [2, 9–13]. The most widespread dinoflagellates species producing PSTs are in the genus *Alexandrium* [14, 15].

PSTs, mainly produced from dinoflagellate and cyanobacteria, is one of the most powerful neurotoxins such as TTX [16]. Currently, the regulation mechanism for PSTs production in cyanobacteria is clear, which mainly refers to 26 functional genes for PSTs biosynthesis [17]. However, the regulation mechanism for PSTs production of harmful dinoflagellates remains poorly understood due to the exceptionally large genome size and high gene copy number of dinoflagellates [18]. Thus, uncovering the factors that influence the biosynthesis of PSTs toxins in dinoflagellates is not only important for understanding PSTs biosynthesis mechanism but also crucial for seafood safety and human health assurance worldwide [19–23].

The environmental factors, such as light, temperature, salinity, nitrogen or phosphorous, have been shown to affect PSTs content for several *Alexandrium* species[24]. For example, while nitrogen limitation causes the toxin content to decrease phosphorous limitation often results in higher toxin cell quotas [25–32].But the salinity and temperature just have limited effects on toxin composition [28, 33].

Bacteria directly or indirectly affect the growth and toxin production of *Alexandrium* species [34–40]. Originally it was considered that the PSTs in dinoflagellate was produced by their associated bacteria, e.g., *protogonyaulax* and *Alexandrium* [37, 41, 42]. However, the subsequent research has shown that the “PSTs” produced by bacteria was actually the analogues of PSTs [43, 44]. The false positive for PSTs detection in bacteria is mainly because of the low concentration. Subsequently many studies indicated that the PSTs in dinoflagellate was produced by dinoflagellates alone, but their associated bacteria could affect their PST production [45–48]. Different bacteria community had different capacity for affecting PSTs production of dinoflagellate [45]. It has been suggested that the effects of bacteria on PSTs production of dinoflagellate can be explained by altered toxin synthesis pathways through algal-bacterium interactions [40]. But the detailed influencing mechanism is still unclear. It is hypothesized that the associated bacteria of dinoflagellate affect the PSTs production by releasing some signals or metabolites that regulate the growth and physiological process of dinoflagellate [45–47]. Thus, understanding the influencing mechanism of associated bacteria for PSTs production of dinoflagellate is not only helpful for underlying the PSTs biosynthesis in dinoflagellate but also useful for controlling the PSTs production by microbial ways.

Yangtze Estuary is the largest estuary in China, which plays an important role for industry, agriculture, aquaculture and marine ecology. However, HABs often happen in Yangtze Estuary due to the environmental pollution and eutrophication, especially for dinoflagellate. Therefore, the production of PST from dinoflagellates, especially from *Alexandrium* species, can cause severe shellfish food poisoning [49]. Due to the difficulty in identifying dinoflagellate by morphological characters, in this study, the eDNA metabarcoding [50] was employed to reveal the community composition of dinoflagellate in Yangtze Estuary. The associated bacteria community diversity of *A. catenella* from Yangtze Estuary was revealed and the functional bacteria strains for affecting PSTs production were screened. Then the physiological process and molecular regulation of *A. catenella* under influence of associated bacteria on PSTs production were further evaluated.

## 2. Materials and methods

### 2.1 Sample collection

*A. catenella* seawater samples collected from Shanghai (*A. catenella*-C, 121.92348°E, 30.862512°N), Shengsi (*A. catenella*-S, 122.454838°E, 30.700638°N) and Zhoushan (*A. catenella*-Z, 122.413229°E, 29.92582°N) in Yangtze Estuary of China were firstly pre-filtered through a 200μm pore-size sieve to remove debris, mesoplankton, and macroplankton. Three subsamples of equal volume from 0.5 m, 1.0 m and 2.0 m depths were collected and mixed as one sample from each plot. Then each water sample (∼500 ml) was subsequently filtered through a 0.22μm pore-size polycarbonate membrane (Millipore, Billerica, MA, USA). The membranes were stored at -80 °C until DNA extraction. The toxicity of *A. catenella* strains from different locations was detected, where the stains with the highest toxicity was selected for experiment analysis.

### 2.2 Dinoflagellate community diversity

The 18S-v4-v5 and 28S primers (Table 1) were used for metabarcoding eukaryotic plankton and dinoflagellates by Illumina-miseq high throughput sequencing. DNA was extracted by OMEGA E.Z.N.ATM Mag-Bind Soil DNA Kit for each sample. PCR amplification of all samples was performed in triplicate. After sequencing, the fastq files were demultiplexed to separate the samples based on the indexes. Reads were then analyzed and assigned using QIIME2, where the PCR primers were removed by cut adapt package and Dada2 was applied for denoising, dereplicate-sequence, filtering chimera and merging the paired reads [51]. Taxonomic classification was performed by feature-classifier module with Naive Bayes [51]. The reference library for ASV classification was referred from Zou et al [52].

**Table 1.**
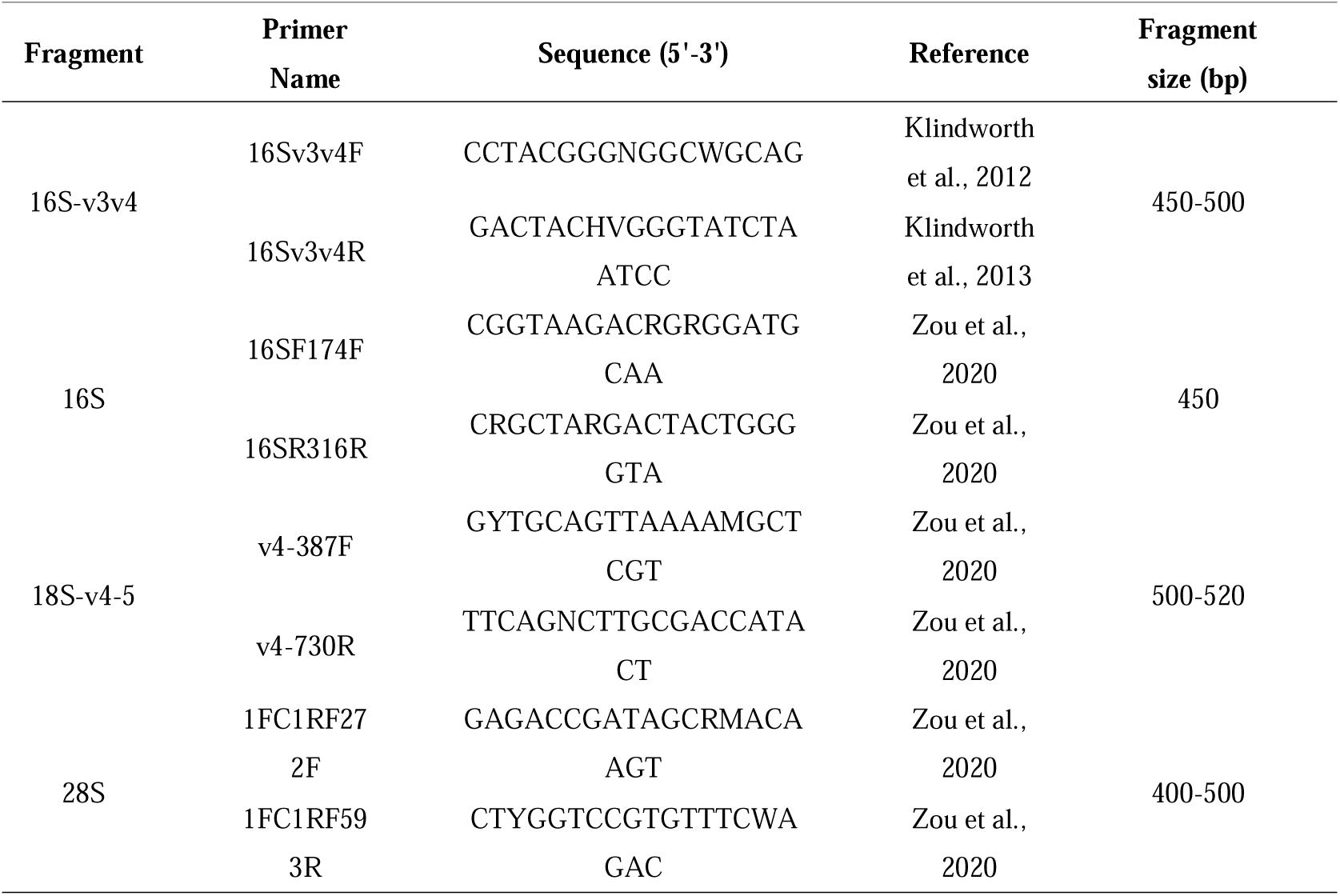
Primers for PCR amplification.

### 2.3 A. catenella isolation, culture and antibiotic treatment

Vegetative cells preliminarily identified as *A. catenella* based on morphological characteristics were isolated by pipette under a microscope (40× zoom), and transferred to a 24-well plates containing culture f/2 medium. These strains were cultured at 19-26°C, 3000-6000lux light and light to dark ratio of 12D: 12N. Then the cultures were expanded from orifices to 250ml triangulated vials and maintained in the laboratory using f/2 medium under the same conditions. The obtained strains were then further detected by 18S sequencing. A set of publicly available 18S sequences of *Alexandrium* downloaded from GenBank was added to the new sequences to build phylogenetic trees by the maximum-likelihood method.

For obtaining axenic strains, cefotaxime sodium and vancomycin hydrochloride were selected as the antibacterial antibiotics after testing with different antibiotic mixtures. 30μg/ml cefotaxime sodium and 50μg/ml vancomycin hydrochloride were added to the sterile f/2 medium for producing axenic *A. catenella* cultures, which was conducted for triple replicates. Tow replicates were used to assess the axenic state of the cultures, while the remainder was used to determine the toxicity. For non-axenic culture with nitrogen deprivation (non-axenic-ndep) the concentration of NaNO3 was set at 1/4 of the control group (2.94 mmol/liter).

### 2.4 Associated bacteria community diversity of A. catenella

The associated bacteria community diversity of *A. catenella* under laboratory culture condition was analyzed by 16Sv3v4 and 16S metabarcoding respectively (Table 1). *A. catenella* culture was filtered with a 0.22μm pore-size polycarbonate membrane to obtain the associated bacteria (Millipore, Billerica, MA, USA). DNA was extracted by OMEGA E.Z.N.ATM Mag-Bind Soil DNA Kit. PCR amplification of all samples were triple. The QIIME2 was used for bioinformatics analysis as above [51].

### 2.5 Isolation and reseeding of associated bacteria

The 2216E medium was used for isolation of associated bacteria of *A. catenella* [53]. Each sample was plated in three plates that were incubated at 20°C, 30°C and 37°C respectively, which were then checked for colonies after 24h, 48h and 72h. Individual bacterial colonies showing different colour, colony size or morphology were re-isolated on the same medium and further characterized morphologically and biochemically with microscope and Gram staining method. Finally, the single clones were further identified by 16Sv3v4 sanger sequencing.

The isolated bacteria were then reseeded to axenic *A. catenella* with antibiotic treatment, through ways of separate strain and different combined strains. While *A. catenella* was cultured to logarithmic phase antibiotic treatment was carried out to get axenic algal solution. The isolated bacterial strains were cultured to logarithmic phase and then added to the algal solution.

### 2.6 Toxicity Type and content test

Taking the algal sap from the logarithmic phase, the algal cells were collected by centrifugation at 6000 rpm·min^-1^ into 50mL polypropylene centrifuge tubes, which were washed and centrifuged several times using ultrapure water to ensure that the culture fluid was thoroughly removed. To the washed algal cells, 20mL of acetic acid solution at a concentration of 0.05 mol/L was added. The algal cells were sonicated under ice bath conditions using an ultrasonic cell disruptor with the power set at 200W and 20s of sonication paused for 40s, for a total of 5min. After sonication, the extraction was continued using ultrasonic assisted extraction for 15 min. the cells were centrifuged at 4000 rpm·min^-1^ for 10 min at 4 °C, and then 1 mL of the supernatant was removed from the centrifuge tube. The supernatant was transferred to an ultrafiltration centrifuge tube (MWCO of 10,000u) and centrifuged at 10,000 rpm·min^-1^ for 10min at 4°C. Finally, the ultrafiltrate was collected into an injection vial and stored at 4°C for subsequent analysis.

The type and content test of PSTs potentially produced by *A. catenella* and associated bacteria was detected by liquid chromatography-mass spectrometry (LC-MS). Samples were analyzed on an Ultimate 3000 ultra-high-pressure liquid chromatography-Q-Exactive electrostatic field Orbitrap high-resolution mass spectrometer system. The column was TSK-gel Amide-80 (3μm, 2mm×15cm). Algal toxin extraction was performed using acetic acid and ultrasound. The instrumental conditions were determined with reference to the method of Zhang et al.[54].

### 2.7 Measurements of growth and biochemical composition

The biomass was detected based on cell number using a hemacytometer (improved double-Neubauer). Soluble sugar was determined by phenol-sulfuric acid method. Soluble proteins were determined using the Thomas Brilliant Blue G-250 staining method. The neutral lipid content was determined by Nile red fluorescence method.

### 2.8 RNA isolation, library construction and transcriptome sequencing

The axenic, non-axenic and non-axenic-ndep *A. catenella* strains were selected for RNA sequencing (RNA-Seq) respectively. Triple replicates of each strain were cultured under the same conditions and were mixed for RNA extraction and sequencing. Total RNA was firstly extracted and qualified by ethanol precipitation protocol and Agilent 2100 bioanalyzer (Thermo Fisher Scientific, MA, USA) respectively. Then the mRNA was purified, fragmented and cDNA produced. The cDNA was subsequently index ligated and enriched. The product was validated on the Agilent Technologies 2100 bioanalyzer for quality control. Finally, the qualified library was sequenced on the Illumina HiSeq platform. Clean reads were mapped to the assembled unique genes by Bowtie2 (v2.2.5) [55] and the expression levels of genes was calculated by RSEM (v1.2.8) and normalized to the FPKM [56].

Functional annotation of genes was achieved by mapping genes to different databases (NT, NR, GO, COG, KEGG, Swiss-Prot, and InterPro) using the software BLAST (v2.2.23) [57]. GO annotation was performed by Blast2GO (v 2.5.0) with NR annotations. PossionDis [58] was used to detect different expression genes (DEGs), and DEGs with a fold change of >2 or <-2 and a false-discovery rate (FDR) of <0.001 were considered significantly differentially expressed genes. GO and KEGG enrichment analysis were performed using Phyper, a function of R. The significant levels of terms and pathways were corrected by Q value with a rigorous threshold (Q <0.05).

## 3. Results

### 3.1 The dinoflagellate community diversity

From the relative abundance of 18S-v4 and 28S ASVs (Figure 1A), it was shown that the Pyrrophyta accounted for a considerable proportion compared to another microalgae phylum, especially higher than Bacillariophyta (Figure 1A). The zooplankton was revealed with a high proportion but the Chlorophyta was uncovered with a low proportion. The dominant dinoflagellate species were mainly included in *Alexandrium*, *Karlodinium*, *Prorocentrum*, Dinophyceae, etc. Thereinto, the *Alexandrium fundyense*, *Alexandrium* sp.*, A. catenella* and *Alexandrium tamarense* dominated high abundance (Figure 1B). The phylogenetic tree showed that the *A. catenella* sequences from Shanghai, Shengsi and Zhoushan (Genbank: OR357631-OR357633) clustered as one clade (Figure 2A). The toxin content of *A. catenella* samples from the three locations was different (Figure 2B), consistent with the difference in bacterioplankton (Figure 2C). The *A. catenella* strain with the highest toxin content from Shengsi was selected for detailed experimental analysis.

**Figure 1.**
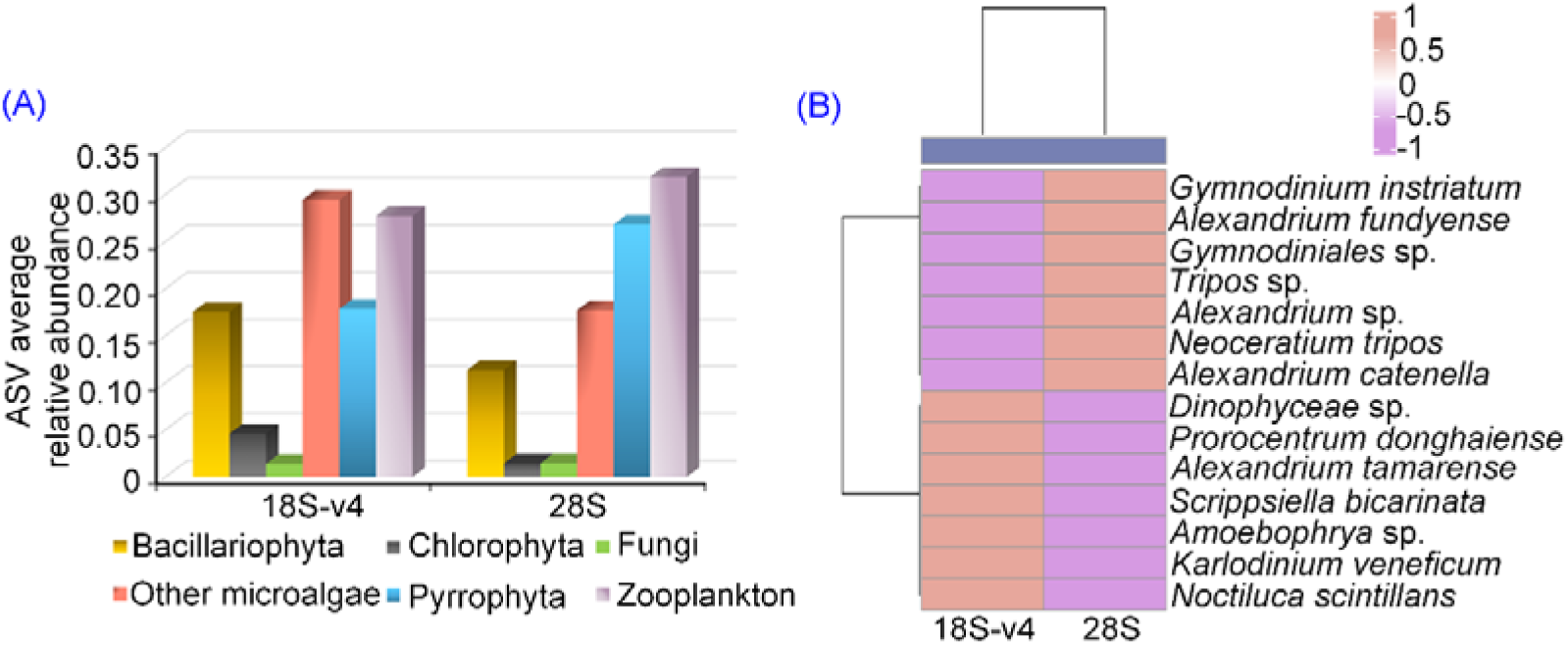
The community diversity of dinoflagellate in Yangtze Estuary. (A): ASV mean relative abundance; (B): Dominant dinoflagellate species.

**Figure 2.**
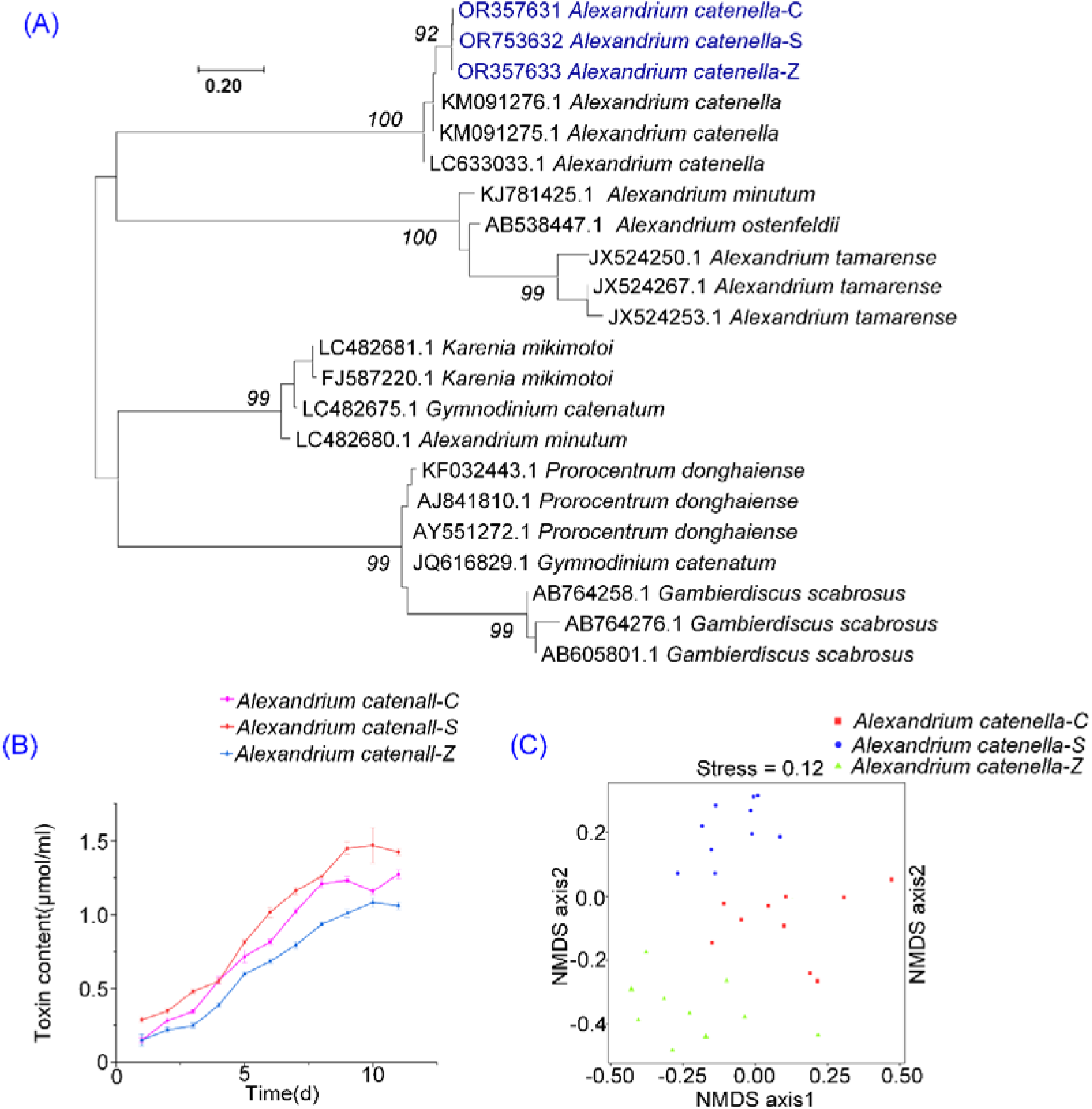
(A): Maximum-likelihood phylogenetic tree of *A. catenella* from 18S. (B): The toxin content of *A. catenella* samples from three locations; (C): The associated bacteria difference of *A. catenella* samples from three locations. The points represent means ± SD of triplicate samples.

### 3.2 The associated bacteria composition of A. catenella

Here the 16S and 16Sv3v4 assignments were combined together to reveal the associated bacteria composition of *A. catenella*. Six phyla, Bacteroidetes, Proteobacteria, Steptophyta, Firmicutes, Bacteroidota and Pseudomonadota, were detected. The dominant classes were Cyanbacteria, Cytophagia, Deltaproteobacteria, Flavobacteriia, Bacilli, Gammaproteobacteria, Alphaproteobacteria and Bacteroidia. The main orders included Alteromonadales, Flavobacteriales, Cytophagales, Rhodobacterales etc. (Figure 3). The dominant associated bacteria species included *Paenibacillus* sp., *Yangia pacifica*, *Formosa* sp., *Bacillus humi*, *Alteromonas macleodii*, *Marinobacter* sp., *Cesiribacter* sp., *Flavobacteriales bacterium* etc. Among them, *Marinobacter* accounted for the highest proportion in 16S and *Paenibacillus* accounted for the highest proportion in 16Sv3v4 (Figure 4).

**Figure 3.**
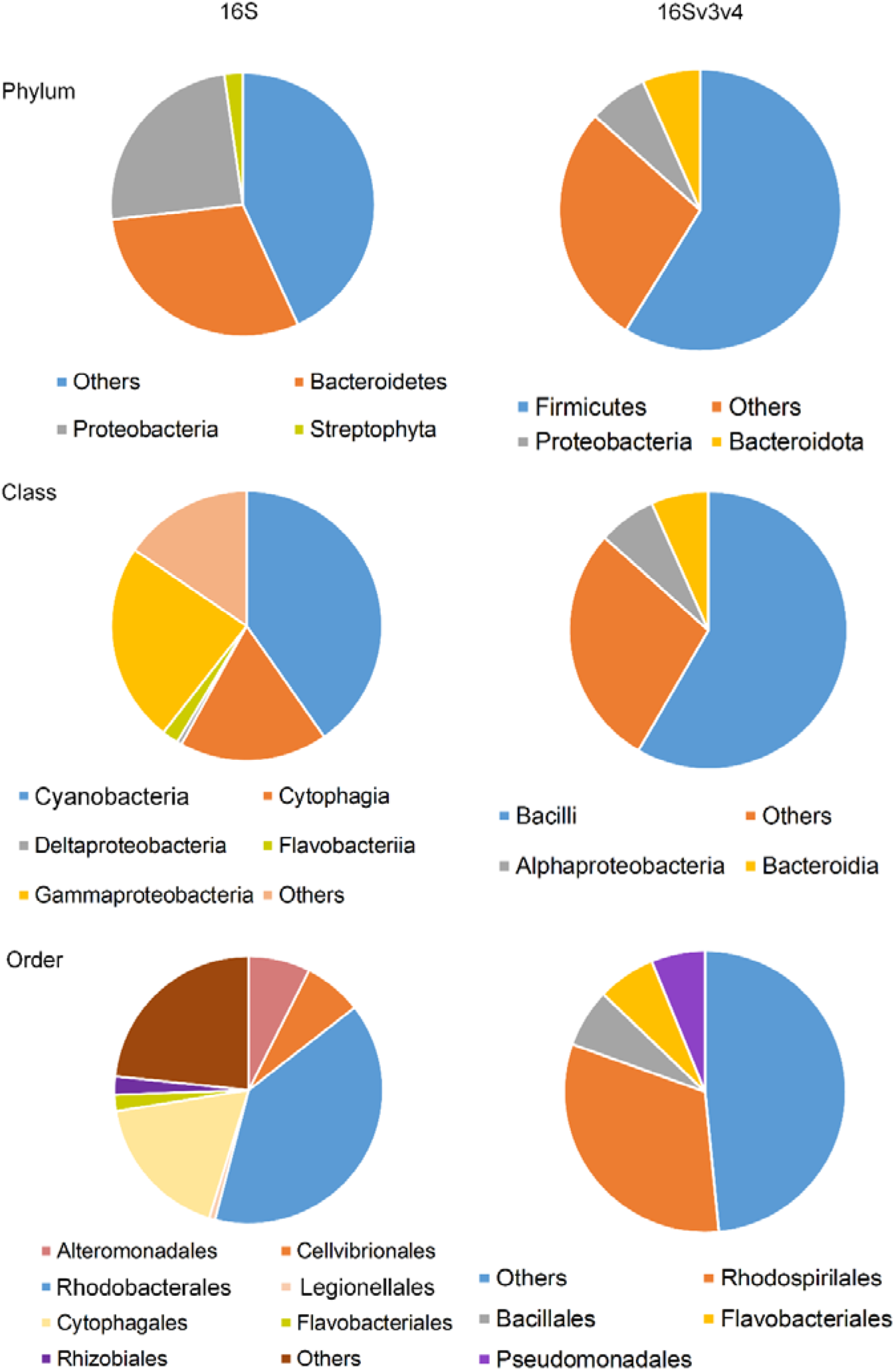
Associated bacteria composition of *A. catenella* from 16S and 16Sv3v4 at phylum, class, order taxonomic level.

**Figure 4.**
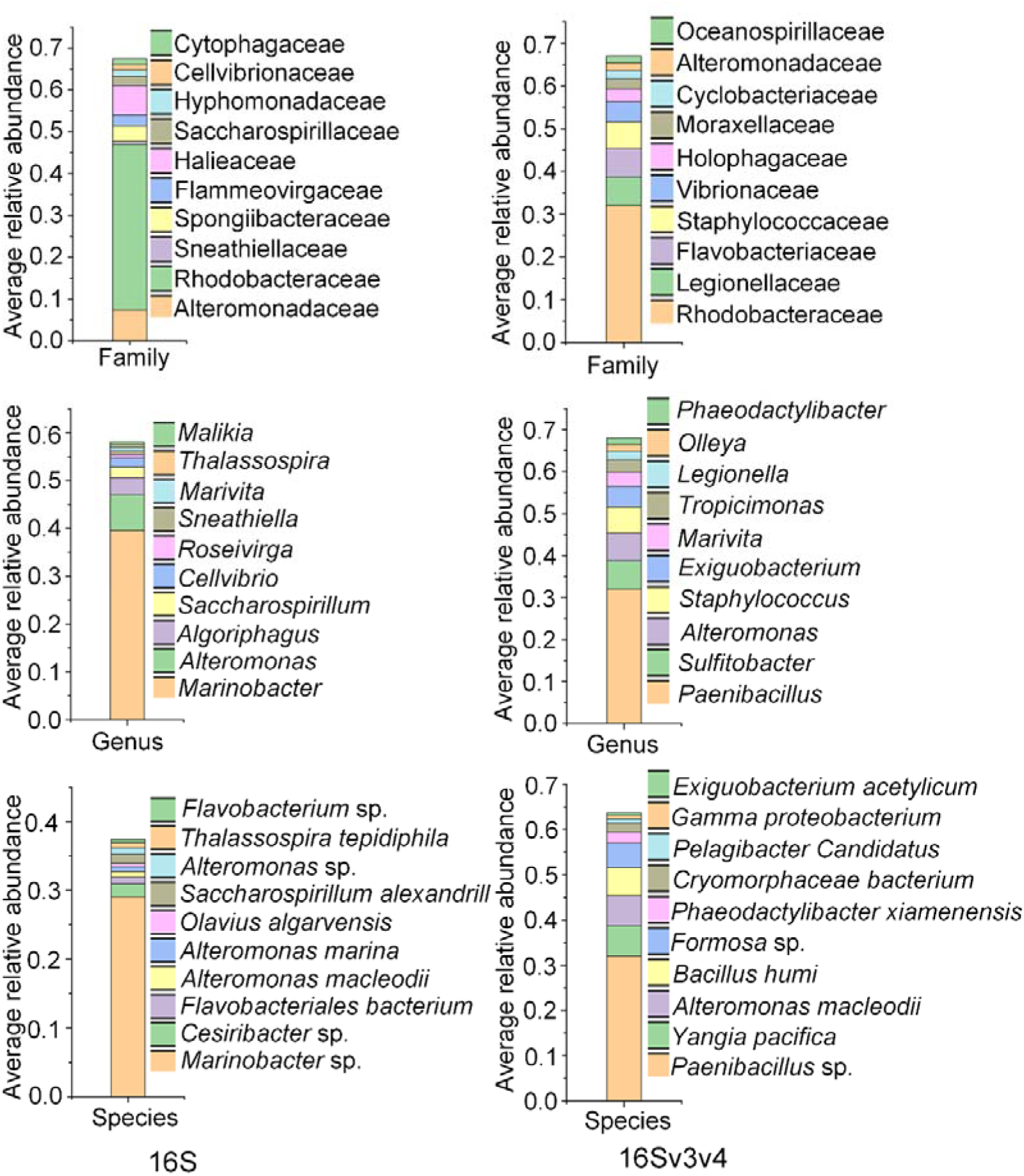
Associated bacteria composition of *A. catenella* from 16S and 16Sv3v4 at family, genus, species taxonomic level.

### 3.3 Associated bacteria isolation and capacity of affecting toxin production test

A total of 31 associated bacteria of *A. catenella* were isolated, which were identified to 13 different bacteria species by 16Sv3v4 sequences (Figure 5A). These were *A. macleodii, Alteromonas gracilis, Alteromonas marina, Thalassospira marina, Alteromonas* sp.*, Exiguobacterium acetylicum, Marinobacter* sp.*, Cellvibrio* sp.*, Spongiibacter* sp.*, Saccharospirillum* sp.*, Saccharospirillum alexandrill, Marinobacter confluentis, Marinobacterium rambilcola* (Genbank: OR958743-OR958746, OR960642-OR960647, OR976523-OR976525). The 13 bacteria were assigned to four classes, six orders and eight families (Figure S1).

**Figure 5.**
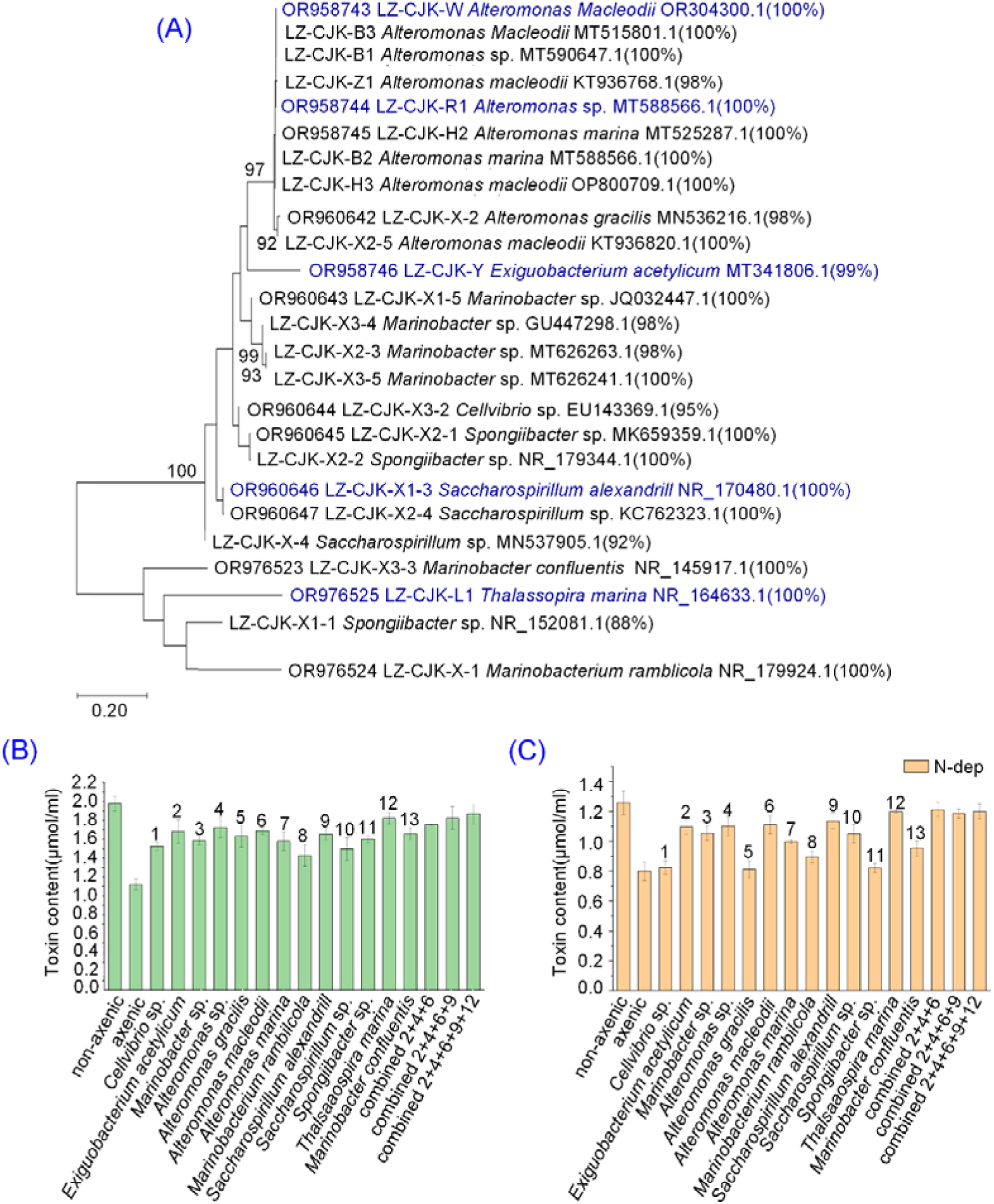
(A): Maximum-likelihood phylogenetic tree of isolated associated bacteria of toxic *A. catenella* strain. (B): The influence of isolated bacteria on toxin production of *A. catenella* under normal culture conditions. (C): The influence of isolated bacteria on toxin production of *A. catenella* under nitrogen deprivation conditions. 1-13 indicates the bacteria serial number for reseeding with *A. catenella*.

With LC-MS analysis, it was showed that no PSTs was detected in the mixed associated bacterial culture since there was no same molecular weight with PSTs component was found (Figure 6A). On the other hand, a molecular weight of 395.0878 was detected in the algal culture, which was the same as GTX toxin at the peak time of 6.475min (Figure 6B, C).

**Figure 6.**
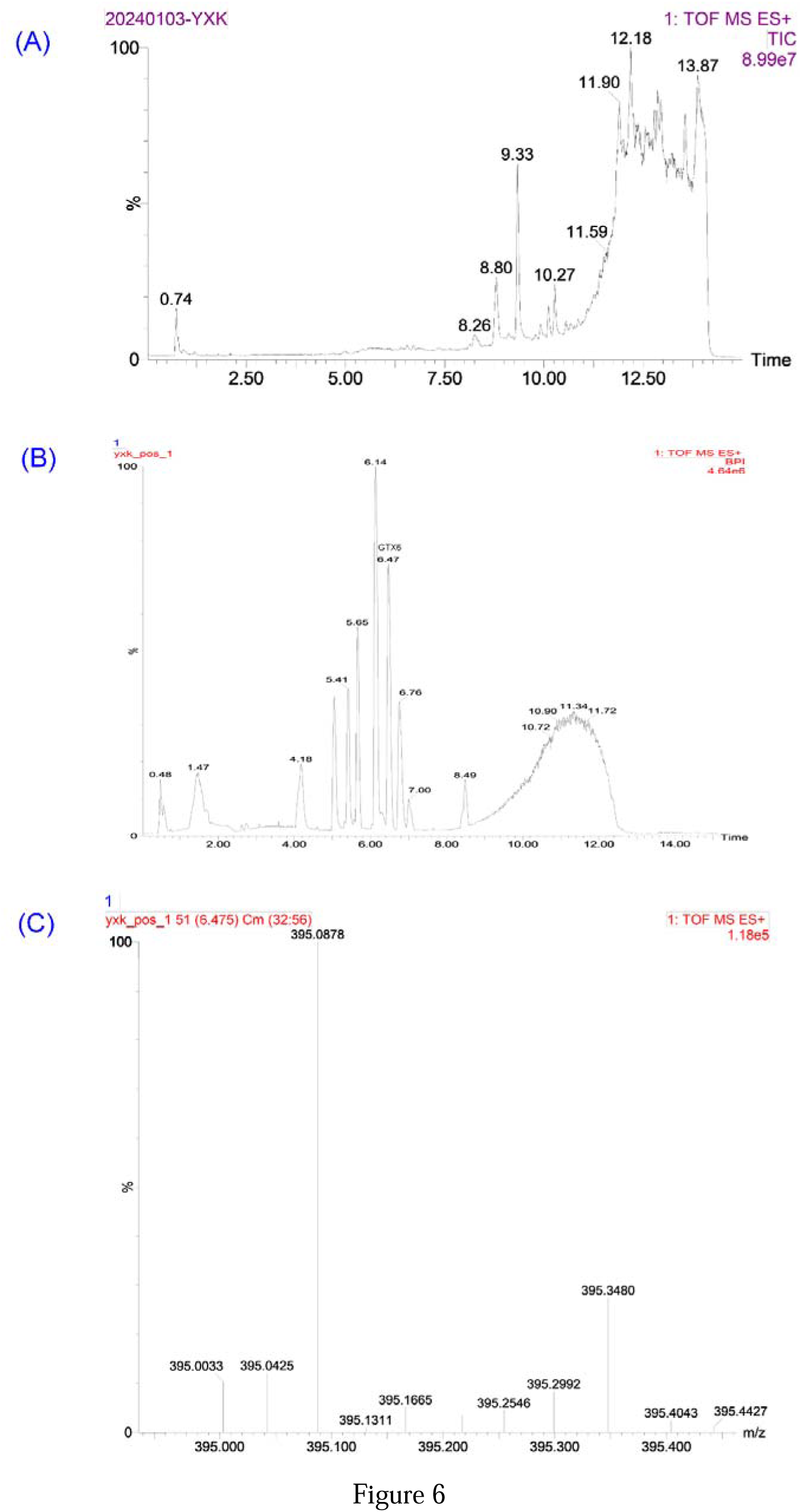
(A): LC-MS mass spectral peaks for mixed bacterial solution. Vertical coordinate is mean relative abundance, horizontal coordinate is time. (B): LC-MS mass spectral peaks of the toxic *A. catenella* strain; (C): The molecular weight of detected GTX toxins (395.0878) from LC-MS peaks of toxic *A. catenella* strain.

The capacity of isolated bacteria for affecting PSTs production of *A. catenella* was tested separately, together with different combined bacteria strains (Figure 5B, C). Under normal cultivation conditions, the non-axenic *A. catenella* strains without antibiotic treatment apparently produced more PSTs than strains with antibiotic treatment (Figure 5B). Generally, the non-axenic *A. catenella* strains with separate and combined isolated bacteria all produced more PSTs than axenic strain, especially for stains with combined bacteria strains (Figure 5B). Moreover, the capacity of isolated bacteria for affecting PSTs production was detected under nitrogen deprivation, to select typical bacteria that affects PSTs production pronouncedly (Figure 5C). It was apparent that the toxin content decreased for almost all treatments under nitrogen deprivation (Figure 5C). Among the 13 bacterial strains, it was showed that *A. catenella* produced more PSTs with five associated bacteria strains, e.g. *E. acetylicum, A.* sp*., A. macleodii, S. alexandrill and T. marina* (Figure 5B, C).

### 3.4 The growth and biochemical composition for axenic and non-axenic A. catenella strains

It was shown that the axenic *A. catenella* strain with antibiotic treatment grew slower than non-axenic *A. catenella* strain without antibiotic treatment during the whole growth cycle under normal cultivation conditions (Figure 7A). Thereinto, the non-axenic *A. catenella* strain with nitrogen deprivation (non-axenic-ndep) grew the slowest (Figure 7A). The non-axenic *A. catenella* strain produced more soluble protein, sugar and neutral lipid than axenic *A. catenella* strain with antibiotic treatment normal cultivation conditions (Figure 7B). On the other hand, the soluble sugar and protein content decreased and the neutral lipid content increased in the non-axenic *A. catenella* strains with nitrogen deprivation (non-axenic-ndep), compared with both axenic and non-axenic *A. catenella* strains under normal cultivation conditions (Figure 7B).

**Figure 7.**
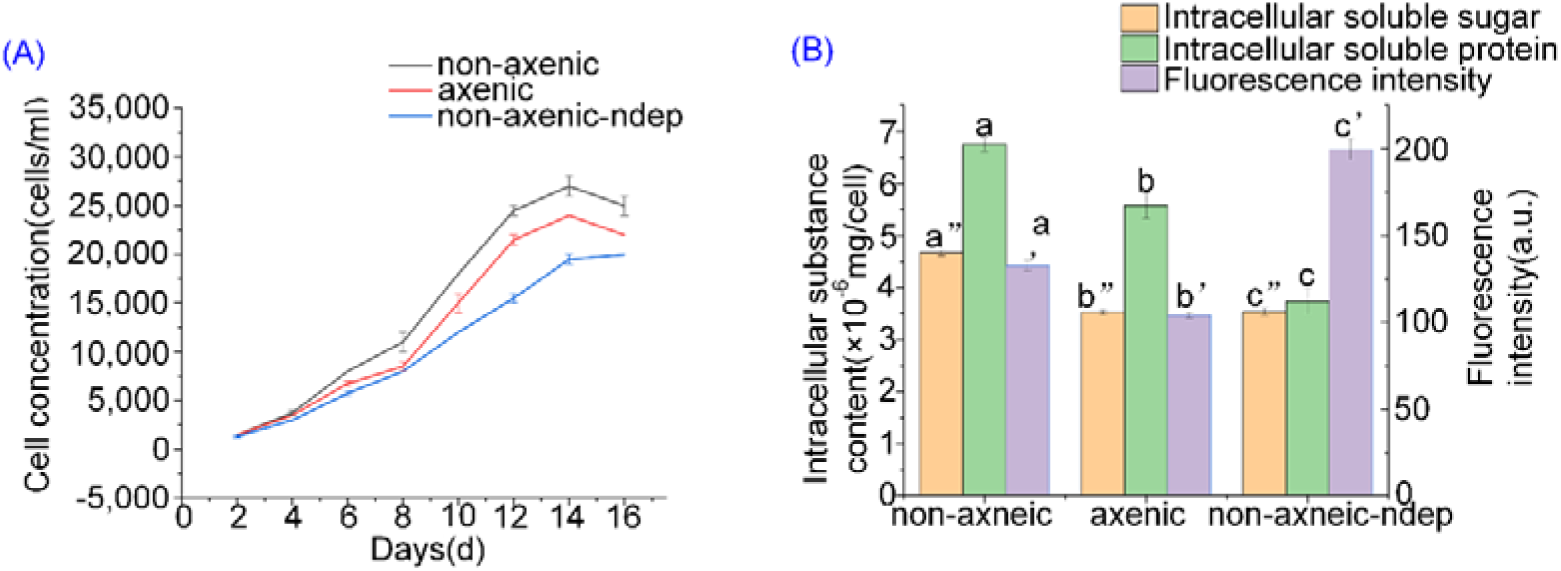
(A): Growth of axenic, non-axenic and non-axenic-ndep *A. catenella* strains; (B): The content of soluble protein, sugar and neutral lipid from non-axenic and non-axenic-ndep *A. catenella* strains. The points represent means **±** SD of triplicate samples. Different letters above the bars indicate significant differences (P < 0.05) between growth condition.

### 3.5 Genetic regulation for PSTs production with influence of associated bacteria

A total of 45,778,990, 48,518,624 and 43,035,798 reads were generated for each treatment of axenic, non-axenic and non-axenic-ndep *A. catenella* strains respectively (Table 2). The clean read ratio of all samples was approximately 97%. The total number of transcripts for three treatments were 145,580, with a mean length 923 bp and 1382 N50 length. A total of 126,626 unigenes were obtained for three treatments, with a mean length of 879 bp and an N50 length of 1327. Thus, the reads and assembly quality showed a good preformation of transcriptome sequencing for functional analysis.

**Table 2.** Sequencing data statistics table.

| Sample | Raw reads | Clean reads | Error rate (%) | Q20 (%) | Q30 (%) | GC content (%) |
| --- | --- | --- | --- | --- | --- | --- |
| non-axenic-ndep | 45778990 | 45067136 | 0.0234 | 98.68 | 95.83 | 56.25 |
| non-axenic | 48518624 | 48028712 | 0.024 | 98.44 | 95.15 | 58.51 |
| axenic | 43035798 | 42337016 | 0.0245 | 98.25 | 94.76 | 56.55 |

Among the annotations of the six databases, the Pfam database showed higher annotation proportion and more unique annotation (Figure 8A). Among the transcription factors, the vast majority of unigenes were assigned to C3H, followed by MYB_superfamily (Figure 8B). The axenic, non-axenic and non-axenic-ndep strains all produced abundant common and distinct unigenes (Figure 8C). The up-regulated and down-regulated genes among three *A. catenella* treatments were different (Figure 8D). The GO functional annotations was performed for all unigenes where the membrane part, cell part, cellular progress, metabolic process, binding and catalytic activity accounted for the top classification categories (Figure S2). The KEGG database assigned the Carbohydrate metabolism, Amino acid metabolism, Translation and Folding, sorting and degradation as the top annotation (Figure S3). It was indicated that the biosynthesis of cofactors was the dominant KEGG pathway annotated between axenic and non-axenic strains (Figure 9A). The dominant DEGs included Glutathione metabolism, Plant-pathogen interaction, Arginine and proline metabolism, Tryptophan metabolism, and spliceosome, etc. (Figure 9B). The PSTs synthesis genes sxtA was found in *A. catenella* strains of all three treatments, which were expressed highly in non-axenic strain (Figure 9B). The main pathways enriched between non-axenic and non-axenic-ndep strains were Phenylpropanoid biosynthesis, Spliceosome, etc. (Figure S4).

**Figure 8.**
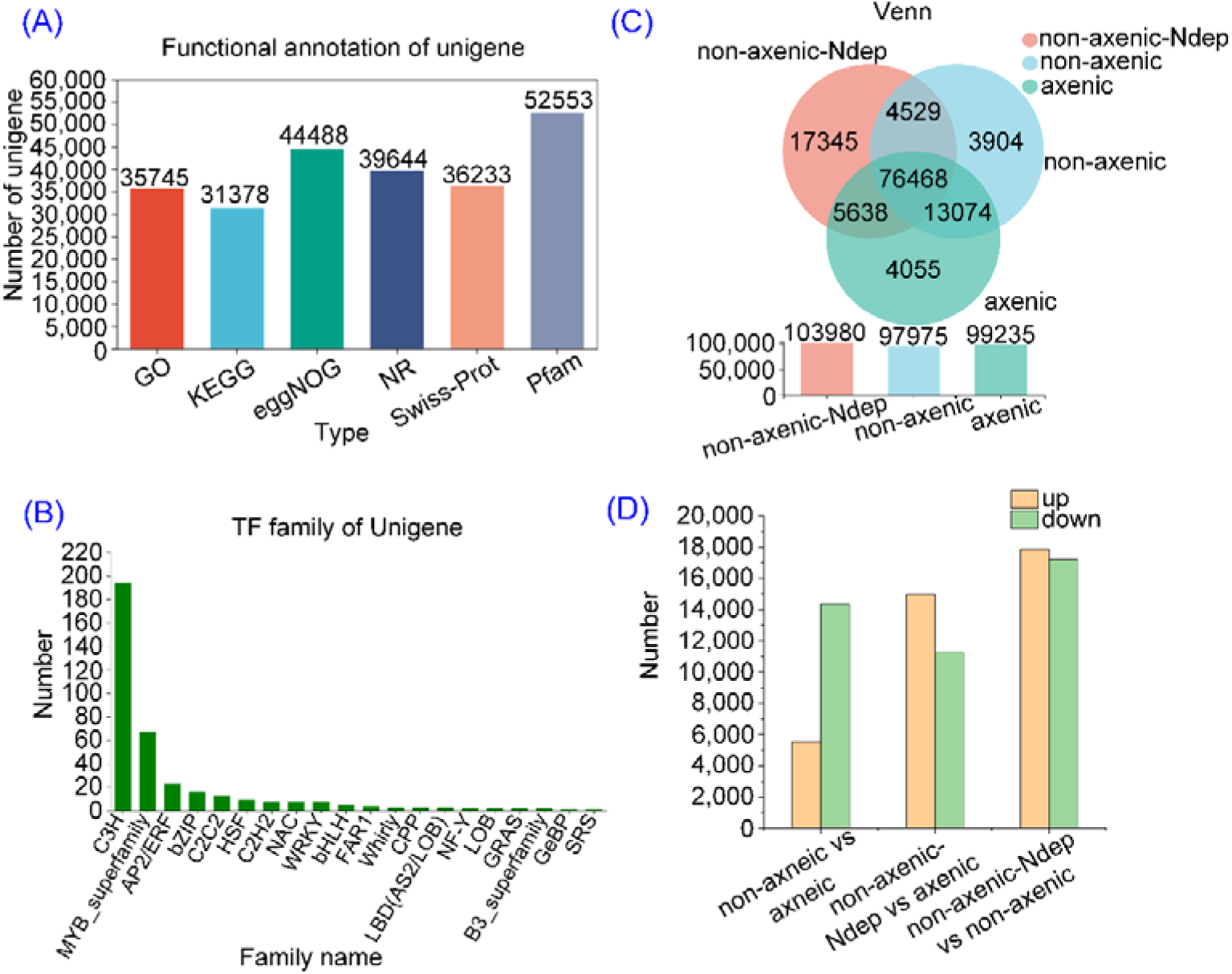
(A): Annotation difference of six databases; (B): Transcription factors annotated from unigenes; (C): Shared and distinct unigenes from axenic, non-axenic and non-axenic-ndep *A. catenella* strains; (D): Up and down-regulated genes among axenic, non-axenic and non-axenic-ndep *A. catenella* strains. For example, non-axenic vs axenic comparison shows that non-axenic strains generated more downregulated genes (green) than upregulated genes (orange) in comparison with axenic strain.

**Figure 9.**
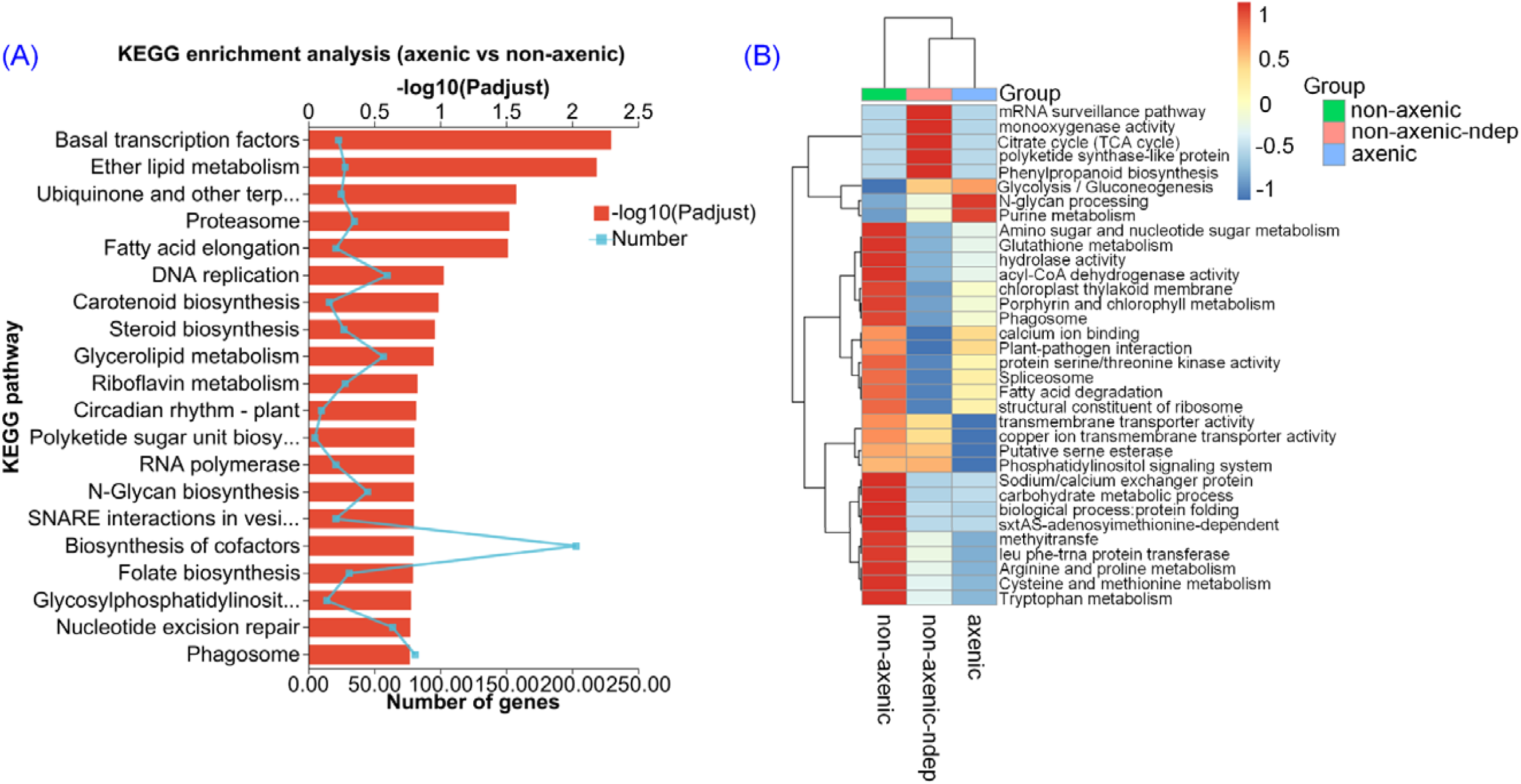
(A): Biosynthesis of cofactors annotated between axenic and non-axenic *A. catenella* strains from KEGG pathway; (B): The dominant different expression genes (DEGs) among axenic, non-axenic and non-axenic-ndep *A. catenella* strains.

The pathway network related with PSTs production was further constructed among axenic, non-axenic and non-axenic-ndep strains, including variable physiological metabolism and PSTs production genes (Figure 10). The pathways Arginine and proline metabolism, Arginine biosynthesis, Fatty acid biosynthesis, TCA cycle and Glutathione metabolism were closely related with sxtA expression, which were all downregulated in axenic strain (Figure 10A). As expected, the plant hormone signal transduction and Indole-3-acetic acid-amido synthetase were up-regulated in non-axenic strain (Figure 10A). Similar with non-axenic vs axenic comparison, generally the non-axenic *A. catenella* strains with nitrogen deprivation produced more downregulated genes than non-axenic *A. catenella* strains under normal cultivation conditions in the various pathways of Glycolysis, TCA cycle, Arginine biosynthesis, Fatty acid biosynthesis, etc. (Figure 10B). Some potential functional genes connecting various pathways related with PSTs production were found, like arginine deiminase and ornithine decarboxylase among Arginine biosynthesis, Arginine and proline metabolism and Glutathione metabolism.

**Figure 10.**
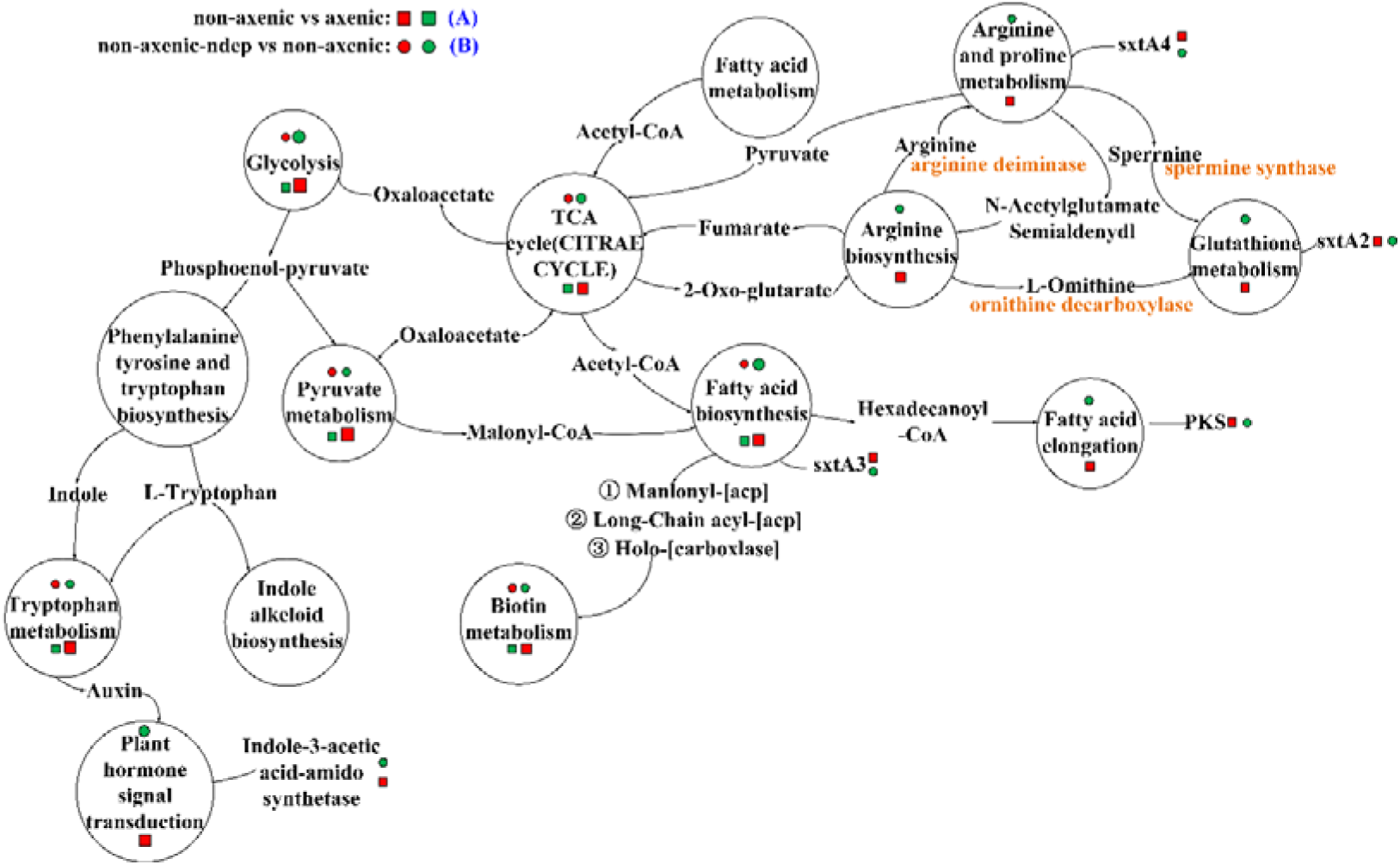
The pathway network related with genetic expression of PSTs production. Red box and circle: upregulated genes, green box and circle: downregulated genes. For example, “non-axenic vs axenic” means the upregulated and downregulated genes non-axenic strains produced compared with axenic strain. The size of box and circle indicates the amount of upregulated and downregulated genes.

## 4. Discussion

The PSTs production by dinoflagellate is associated with bacteria [59, 60]. The PSTs control by microbes is becoming important since the microbes play a key role in microalgae’ growth, metabolism and toxin production [61–63]. Elucidating the associative mechanism between dinoflagellate and associated bacteria can control the PSTs production of dinoflagellate in environmental ways.

The metabarcoding results here indicated that the dinoflagellate and *Alexandrium* were dominant microalgae phylum and species in Yangtze Estuary, suggesting that the PSTs produced by dinoflagellate in China coast is becoming serious. Actually, the dominant phytoplankton in Yangtze Estuary has been transferred from diatom to dinoflagellates in recent years [64]. The toxin-producing capacity and associated bacteria community diversity of *A. catenella* from three Yangtze Estuary locations both differed from each other, indicating that the associated bacteria possibly affect the PSTs production of *A. catenella*. Green et al. (2010) also found that the associated bacteria diversity of *G. catenatum* was related with environmental factors and speculated that the associated bacteria regulated the physiological ecological progress to affect PSTs production.

The LC-MS detection showed that the PSTs produced by *A. catenella* was GTX and the associated bacteria did not produce any component of PSTs. This is consistent with previous reports that the associated bacteria did not produce PSTs and the “PSTs” detected from bacteria may be the analogues of PSTs or false positive [43, 44]. Apparently, the PSTs production capacity of axenic *A. catenella* strains with antibiotic treatment decreased compared with non-axenic strains, which is consistent with previous reports that the associated bacteria affected the PSTs production of dinoflagellate [45–48].

The capacity of associated bacteria for affecting PSTs production was further detected. The *A. catenella* stains co-cultured with separate and combined associated bacteria both produced higher levels of PSTs than axenic *A. catenella* stains, especially with combined associated bacteria. Albinsson et al. [45] also demonstrated that different bacterial communities had different effects on *G.catenatum* toxin production. These further confirm that the associated bacteria affect the PSTs production of dinoflagellate. In this case, it is important to untangle the regulation mechanism of bacteria for PSTs production of dinoflagellate.

The axenic *A. catenella* strains grew slowly compared with non-axenic strains, indicating that the associated bacteria also affect the growth of dinoflagellate. Likewise, the bacterial associates can modify growth dynamics of *Gymnodinium catenatum* [65]. These support the concept that algal–bacterial interaction is an important structuring mechanism in phytoplankton communities. Also the contents of soluble sugar, soluble protein and neutral lipid in non-axenic *A. catenella* strains were somewhat higher than that in axenic strains. All these suggest that the associated bacteria affect the PSTs production of dinoflagellate by regulating growth and physiological metabolism [38–40]. On the other hand, the PST production capacity of non-axenic *A. catenella* strains under nitrogen deprivation were both decreased compared to axenic and non-axenic strains under normal culture conditions, indicating that nitrogen is an important factor for toxin production of dinoflagellate since it is the main component of PSTs [28–30, 32].

The sxt gene cluster for STX synthesis has been elucidated in cyanobacteria [17, 66–68]. Among the 26 sxt genes revealed in cyanobacteria, only several sxt genes were found in dinoflagellate, including sxtA, sxtG and sxtO [69]. The biosynthesis of PSTs starts with sxtA [66, 69]. However, the functional mechanism of sxt for PSTs production is still unclear in dinoflagellate. The polyketide synthase-like (PKS) is another important catalyzing enzyme for carbon skeleton formation of algae toxins [70]. Here, both sxtA and PKS were both included in the DEGs. We found that the sxtA expression was related with TCA cycle, Arginine biosynthesis, proline metabolism, Glutathione metabolism, and Fatty acid biosynthesis, which were all downregulated in axenic strains. Also, Arginine is the main elements of PSTs. Thus, here it is clear that the associated bacteria enhance PSTs production of *A. catenella* by upregulating the expression of sxtA genes and their associated physiological pathways. Other sxt genes, such as sxtG and sxtO, were not detected in this study, which may be related with the intraspecific difference that different strains within one species have different toxicity and possible regulation mechanism [59, 71]. Consistent with the physiological analysis, the genetic expression related with PSTs production was generally downregulated in *A. catenella* strain with nitrogen deprivation, confirming the nitrogen importance in molecular regulation.

Compared with axenic strain, the Tryptophan metabolism and Plant hormone signal transduction were also upregulated in non-axenic *A. catenella* strain, consistent with their growth tendency. This indicates that the associated bacteria can also promote the growth of *A. catenella* except the toxin production. It has already been reported that the associated bacteria advanced the growth of diatom by releasing signal molecules [72, 73]. Finally, some functional genes that were possibly related with the PSTs production were revealed. So, the future studies will focus on uncovering the determinant signal molecular released by associated bacteria and their related regulation pathways for affecting PSTs production of dinoflagellates.

## 5. Conclusion

The toxin production capacity, physiological process and molecular regulation among *A. catenella* strains of axenic, non-axenic and non-axenic with nitrogen deprivation were compared. Thirteen associated bacteria for affecting PSTs production of *A. catenella* were selected. Compared with axenic strain with antibiotic treatment, the non-axenic strain with associated bacteria produced more PSTs contents, soluble sugar, soluble protein and neutral lipid. The biosynthesis of cofactors and spliceosome were the dominant pathways between axenic and non-axenic strains. The pathways Arginine and proline metabolism, Arginine biosynthesis, Fatty acid biosynthesis, TCA cycle and Glutathione metabolism were closely related with sxtA expression. The nitrogen plays key role in PSTs production of *A. catenella* through affecting physiological and molecular regulation. These findings highlight the importance of associated bacteria in affecting PSTs production of dinoflagellate through physiological and molecular regulation.

## Credit authorship contribution statement

Shanmei Zou conceived the design and wrote the manuscript. Xinke Yu conducted the experiments. Shanmei Zou, Xinke Yu, Tiantian Sun analyzed the data. Xinke Yu and Xuemin Wu collected the samples.

## Data availability

The NGS sequence data of all samples obtained in this study are deposited in NCBI. The SRA accession for the NGS sequences is PRJNA992270, PRJNA998555. The submission information for each sample mainly includes the reads sequence in fastq form, the sampling location, the sequence platform.

## Acknowledgement

The bioinformatics analysis of the project was supported by the Bioinformatics Center of Nanjing Agricultural University.

## Figure legends

**Figure S1.**
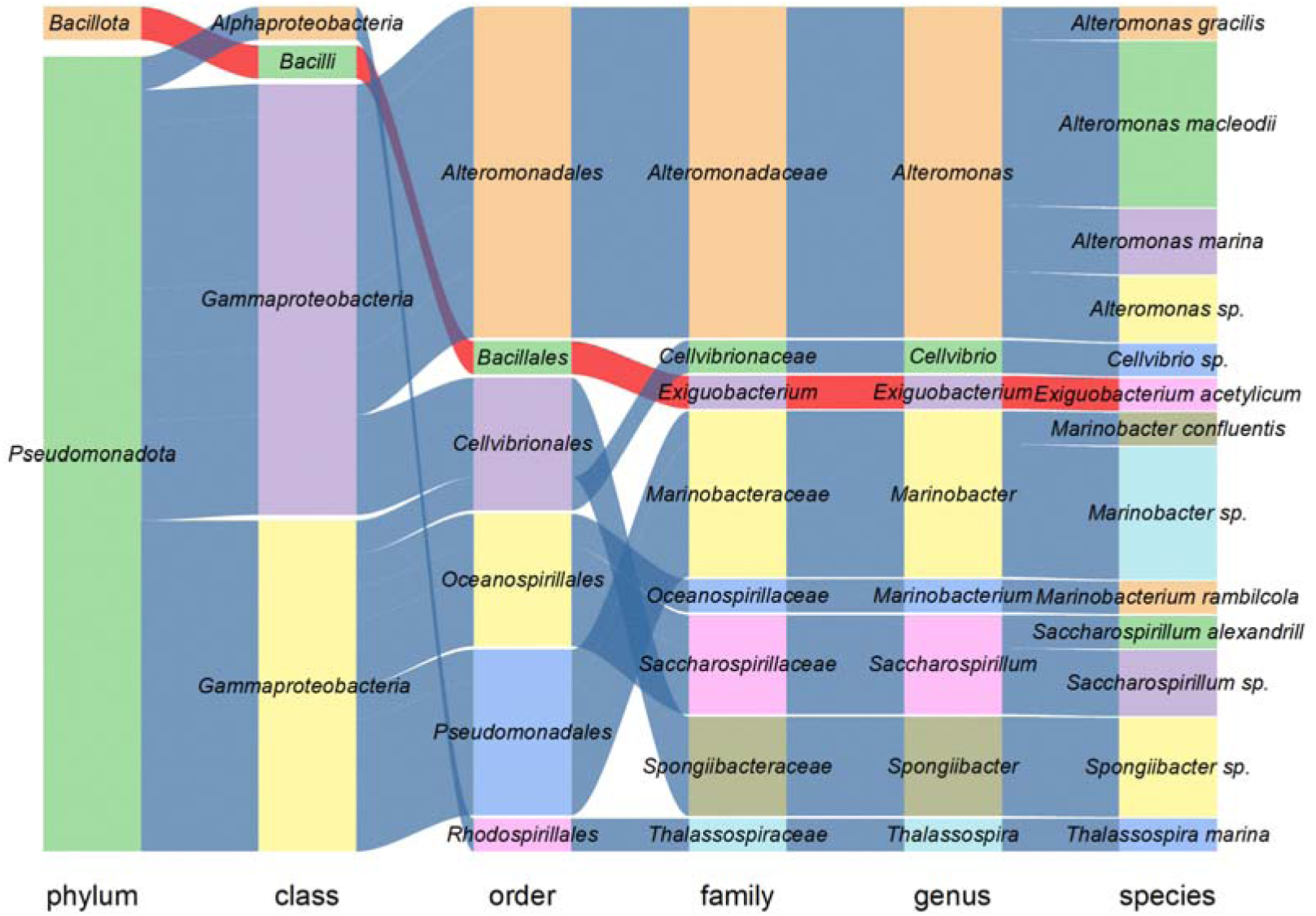
Assignments of isolated associated bacteria at each taxonomic level.

**Figure S2.**
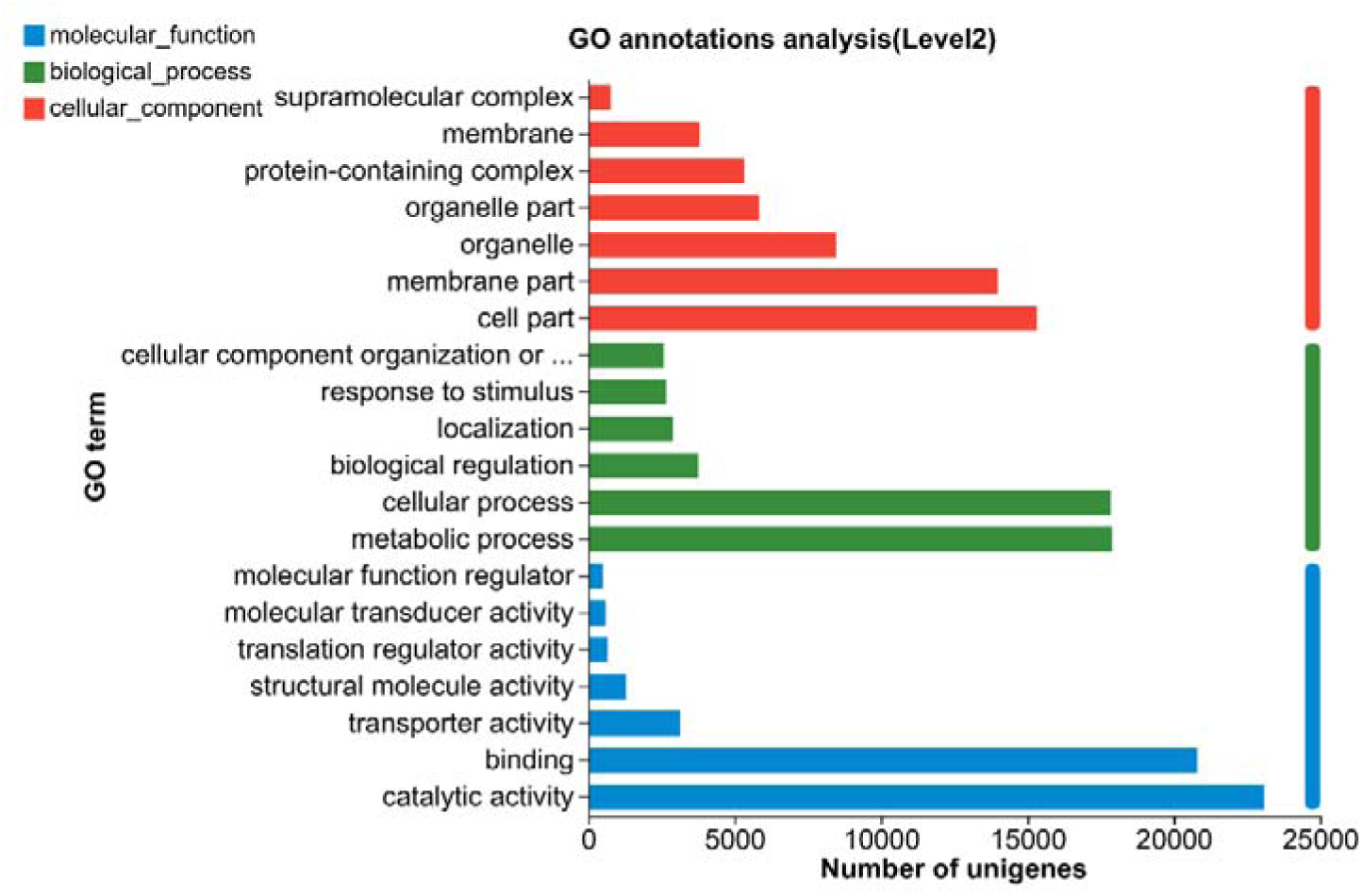
The GO functional annotation performed for all unigenes.

**Figure S3.**
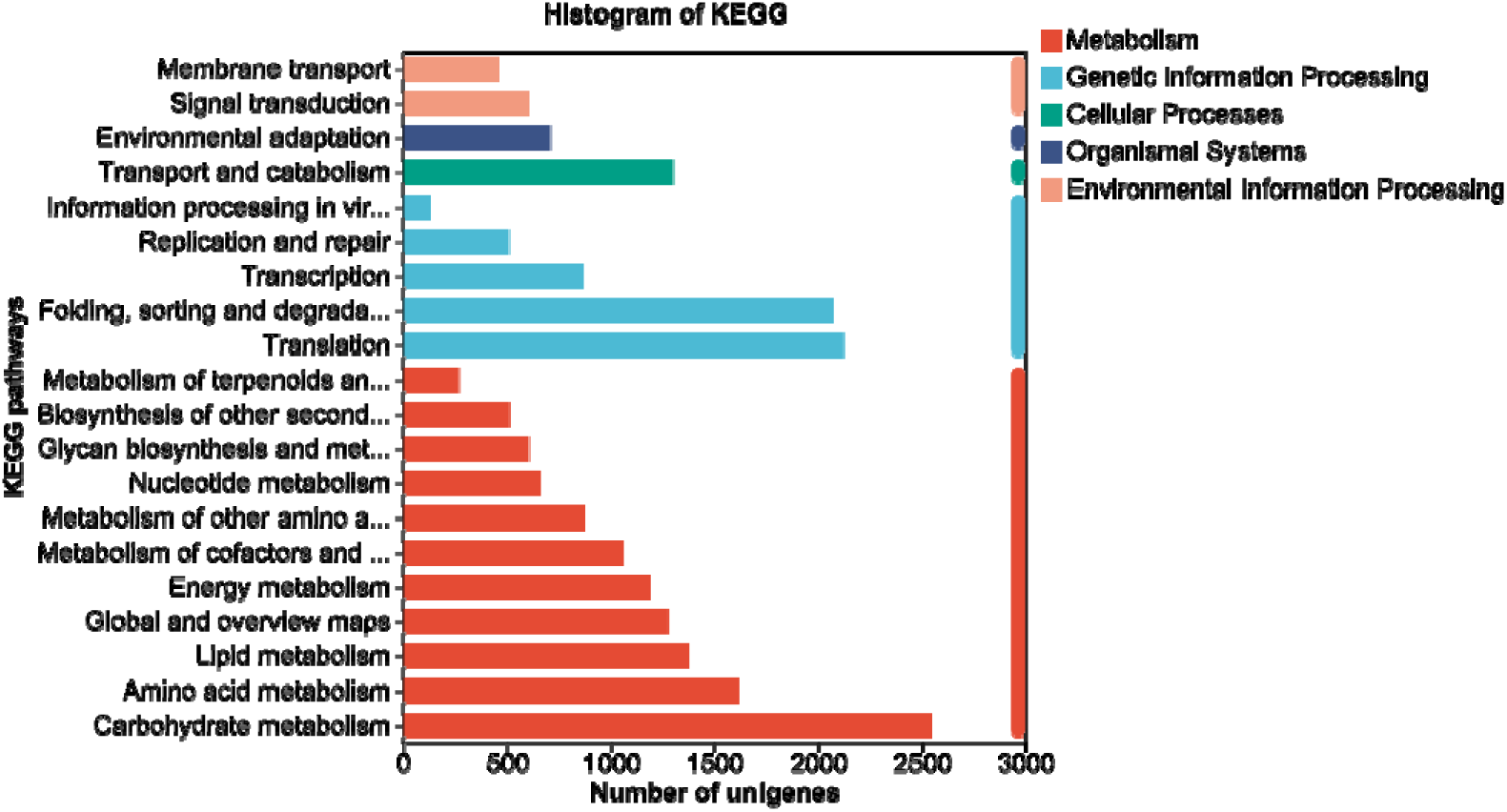
The KEGG pathways assigned.

**Figure S4.**
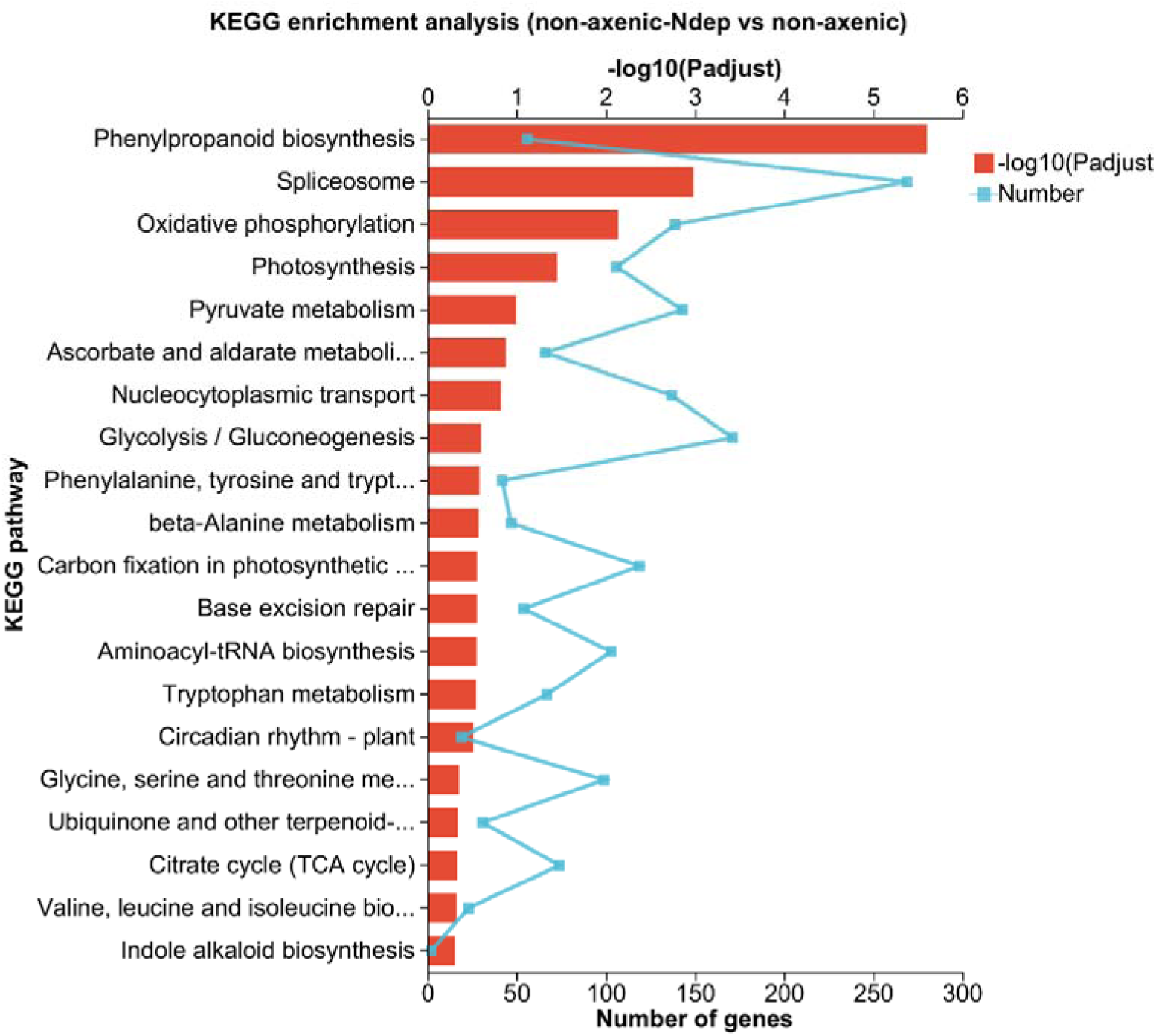
The main pathways enriched between non-axenic and non-axenic-ndep *A. catenella* strains.

